# Osteoarthritis: articular chondrocyte inflammatory signaling leads to enhanced gene expression and function of mechanotransduction channel Piezo1 as a pathogenic feed-forward mechanism

**DOI:** 10.1101/2020.05.23.112565

**Authors:** Whasil Lee, Robert J. Nims, Alireza Savadipour, Holly Leddy, Fang Liu, Amy McNulty, Yong Chen, Farshid Guilak, Wolfgang B. Liedtke

**Author notes:** contributed equally (Farshid Guilak, Wolfgang B. Liedtke). University of Rochester, Biomedical Engineering and Dept. of Pharmacology and Physiology, Rochester, NY, USA.

## Abstract

Osteoarthritis (OA) is a painful and debilitating condition of synovial joints without any disease-modifying therapies (1, 2). We previously identified mechanosensitive PIEZO channels, PIEZO1 and PIEZO2, both expressed in articular cartilage, to function in chondrocyte mechanotransduction in response to injury (3, 4). We therefore asked whether interleukin-1-mediated inflammatory signaling, as occurs in OA, influences Piezo gene expression and channel function, thus indicative of maladaptive reprogramming that can be rationally targeted. Primary porcine chondrocyte culture and human osteoarthritic cartilage tissue were studied.

We found that interleukin-1α (IL-1α) upregulated Piezo1 in porcine chondrocytes. Piezo1 expression was significantly increased in human osteoarthritic cartilage. Increased Piezo1 expression in chondrocytes resulted in a feed-forward pathomechanism whereby increased function of Piezo1 induced excess intracellular Ca^2+^, at baseline and in response to mechanical deformation. Elevated resting state Ca^2+^ in turn rarefied the F-actin cytoskeleton and amplified mechanically-induced deformation-microtrauma. As intracellular substrates of this novel OA-related inflammatory pathomechanism, in porcine articular chondrocytes exposed to IL-1α we discovered that enhanced Piezo1 expression depended on p38 MAP-kinase and transcription factors HNF4 and ATF2/CREBP1. CREBP1 directly bound to the proximal *PIEZO1* gene promoter.

In ensemble, these signaling and genetic re-programming events represent a novel and detrimental Ca^2+^-driven feed-forward mechanism that can be rationally targeted to stem the progression of OA.

**Significance Statement:** Osteoarthritis affecting weight-bearing joints is a global health problem, causing loss of mobility and enormous healthcare costs. Disease-modifying approaches are lacking. Here, we report a new cellular mechanism of inflammatory signaling in chondrocytes, the cellular substrate of cartilage. We show how osteoarthritis-relevant levels of interleukin-1α reprogram articular chondrocytes so that they become more susceptible to mechanical trauma, which chondrocytes sense via Piezo1/2 mechanosensitive ion channels. We uncover that IL-1α enhances gene expression of Piezo1 in primary articular chondrocytes underlying Piezo1 gain-of-function. We elucidate the new signaling pathway, from membrane to nucleus, including transcription factors that enhance Piezo1-expression. We also define detrimental effects of gain-of-function of Piezo1, for mechanotransduction and at-rest, that suggest this new reprogramming mechanism to contribute to osteoarthritis pathogenesis.

## Introduction

OA is a significant global health issue with increasing population age as well as rising obesity rates (5-7). OA is characterized by progressive joint degeneration and pain, leading to significant disability and lack of mobility that further aggravates other age-associated conditions. Due to the multifactorial etiology of the disease and the lack of a full understanding of OA pathogenesis, there are no disease modifying OA drugs (DMOADs) currently available (8, 9). However, growing evidence has documented increased levels of interleukin-1 (IL-1) based inflammatory signaling in chondrocytes, the sole cell population in healthy articular cartilage (10-14). Articular chondrocytes express functional IL-1-receptor and respond to both isoforms of IL-1 (α and β) potently, through catabolic and anti-anabolic activities (15). Mechanical factors, over protracted time multiple iterative microtrauma, play a critical role in OA pathogenesis through alterations in cell-mediated mechanotransduction in cartilage (6, 16-21), and may interact with injurious loading to enhance cartilage degeneration (22). At the molecular level, we have described the presence of both mechanosensory Piezo ion channels (PIEZO1 and PIEZO2) in chondrocytes, which function synergistically in response to injurious mechanical loading (3). In the present study, we address the unanswered question whether joint inflammation, as occurs in OA, affects gene regulation and function of Piezo ion channels as a pro-pathogenic OA mechanism. We provide affirmative and mechanistic answers on how IL-1-mediated inflammatory signaling in articular chondrocytes upregulates *PIEZO1* gene expression and function.

## Results

### OA-relevant levels of pro-inflammatory IL-1α enhance expression of *PIEZO1*

First, we found *PIEZO1* mRNA significantly increased in porcine primary articular chondrocytes in response to IL-1α, over a range of physiologically and pathologically-relevant concentrations (Fig. 1A). Dose-dependence was clearly present between control, 0.1 ng and 1 ng IL-1α, followed by a plateau of *PIEZO1* mRNA levels when comparing 1 ng and 10 ng IL-1α. This increase also manifested as elevated PIEZO1 protein expression as shown by up-regulated PIEZO1 immunolabeling of primary porcine chondrocytes (Fig. 1B), using a PIEZO1 antibody that we verified in a human *PIEZO1*^-/-^ cell line (Suppl. Fig. 1). Expression of *PIEZO2* was increased in response to IL-1α, but not to significant degree, while expression of *TRPV4*, a well-characterized chondrocyte mechanosensor (23-26), was not regulated by IL-1α (Suppl. Fig. 2). We also measured *PIEZO1* expression in human cartilage, where we detected significantly elevated PIEZO1 protein by immunolabeling in osteoarthritic cartilage compared to normal controls (Fig. 1C) again using the validated PIEZO1 antibody. Thus, IL-1α-signaling, at OA-relevant concentrations, evokes significantly increased *PIEZO1* mRNA expression, a finding confirmed at the protein level in articular chondrocytes and human osteoarthritic cartilage lesions. We therefore decided to test PIEZO1 function by use of Yoda-1, a specific PIEZO1 activator, and Ca^2+^-imaging (27). With Yoda-1, primary porcine chondrocytes pretreated with IL-1α exhibited enhanced Ca^2+^ signaling, with a robustly accelerated signal increase and a resulting vastly increased amount of Ca^2+^ entering the cell (Fig. 1D, Suppl. Fig. 3). We conclude that IL-1α inflammatory signaling of articular chondrocytes increases PIEZO1 expression and function in chondrocytes. Building on this finding, the concept emerges that inflammation deleteriously predisposes chondrocytes to become more sensitive to injurious levels of loading. Our finding that *TRPV4*, a mechanosensor that transduces physiologic levels of loading in chondrocytes, is not regulated by IL-1α suggests inflammation selectively alters chondrocyte mechanotransduction pathways. Based on this result, we conducted more in-depth Ca^2+^-signaling studies in response to controlled single-cell mechanical stimulation with or without IL-1α-mediated inflammation.

**Fig. 1.**
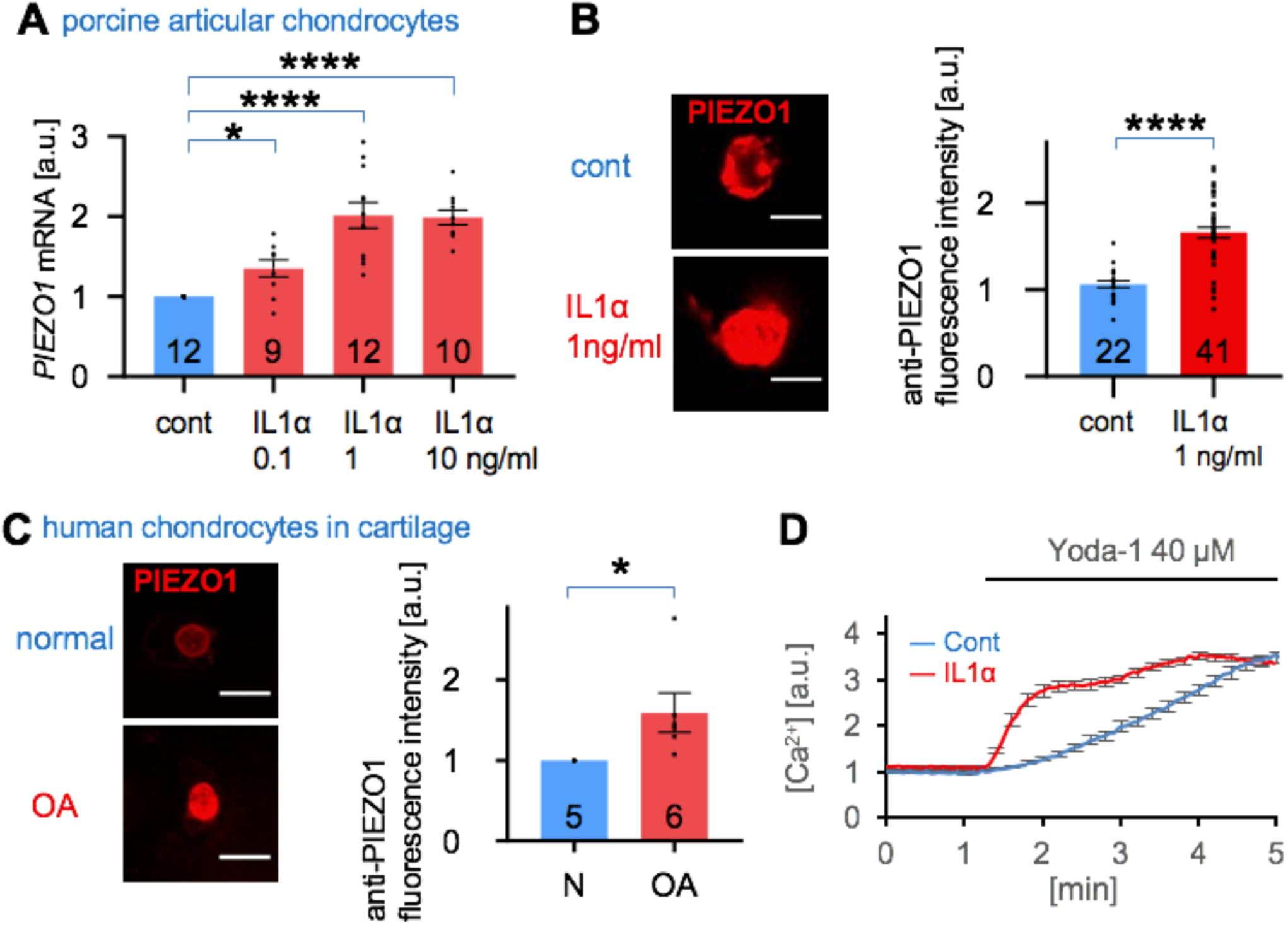
Increased Piezo1 expression in IL-1α-treated porcine chondrocytes, and human osteoarthritic cartilage. **(A)** *PIEZO1* mRNA level in control and IL-1α-treated porcine chondrocytes. IL-1α increases *PIEZO1* mRNA levels; number of independent experiments, primary cells from a separate joint, indicated in bars; a.u. – arbitrary units. **(B)** (Left) Representative confocal micrographs of IL-1α-treated porcine chondrocytes, immunolabeled for PIEZO1. (Right-hand bar diagram) Significantly increased *PIEZO1* protein expression is found in IL-1α-treated porcine chondrocytes vs control. **(C)** (Left) Representative confocal micrographs of healthy and osteoarthritic human chondrocytes; PIEZO1-specific immunolabeling of chondrocytes in human cartilage tissue (see also Suppl. Fig 2 for antibody validation using *PIEZO1*^-/-^ cells), scale bar = 10μm. (Right) Relative expression level PIEZO1 in healthy and OA chondrocytes, average shown for n=5∼6 human cartilage samples, n≥166 cells per group. **(D)** IL-1α-treated porcine chondrocytes show different Ca^2+^ dynamics in response to application of the PIEZO1-selective activating small molecule, Yoda-1. Note accelerated onset, then plateau in the IL-1α-treated cells vs protracted influx in control chondrocytes, indicative of a gain-of-function of PIEZO1-mediated Ca^2+^ signaling under inflammatory conditions, see also Suppl. Fig. 3 for analysis of area-under-the-curve. Bars represent mean±S.E.M; for group comparison: t-test for B, C; 1W-ANOVA, post-hoc Tukey test for A; * p < 0.05, ****p<0.0001 significantly different from control.

### Piezo1 gain-of-function elevates baseline [Ca^2+^]_i_ and renders chondrocytes mechanically hypersensitive

Isolated primary porcine chondrocytes were subjected to recording their steady-state intracellular calcium levels, [Ca^2+^]_i_, using ratiometric Ca^2+^-imaging, in the presence or absence of IL-1α. Chondrocyte [Ca^2+^]_i_ increased 37% in the presence of 0.1 ng/ml IL-1α. However, no further increase in [Ca^2+^]_i_ was observed with higher concentration of IL-1α (Fig. 2A-B). We conclude that IL-1α exposure, at 0.1 ng/mL, profoundly changes articular chondrocytes’ functional state by increasing basal steady-state [Ca^2+^]_i_ by >30%. We next conducted controlled compression of isolated chondrocytes, using atomic-force microscopy (AFM), with a tip-less cantilever to exert dynamic loading of 300 nN at a rate of 1 µm/sec ramp-speed and ramps every 10 sec, while simultaneously recording [Ca^2+^]_i_ (Fig. 2C). The Ca^2+^-response to this form of mechanical load was also significantly amplified with exposure to IL-1α, again showing an appreciable response to IL-1α at 0.1 ng/mL (≈80% increase), with no further increase at 1 and 10 ng/mL (Fig. 2D) (28, 29). Thus, IL-1α as an inflammatory signal makes articular chondrocytes more sensitive to dynamic mechanical compression at a high but physiologically-relevant force of 300 nN.

**Fig. 2.**
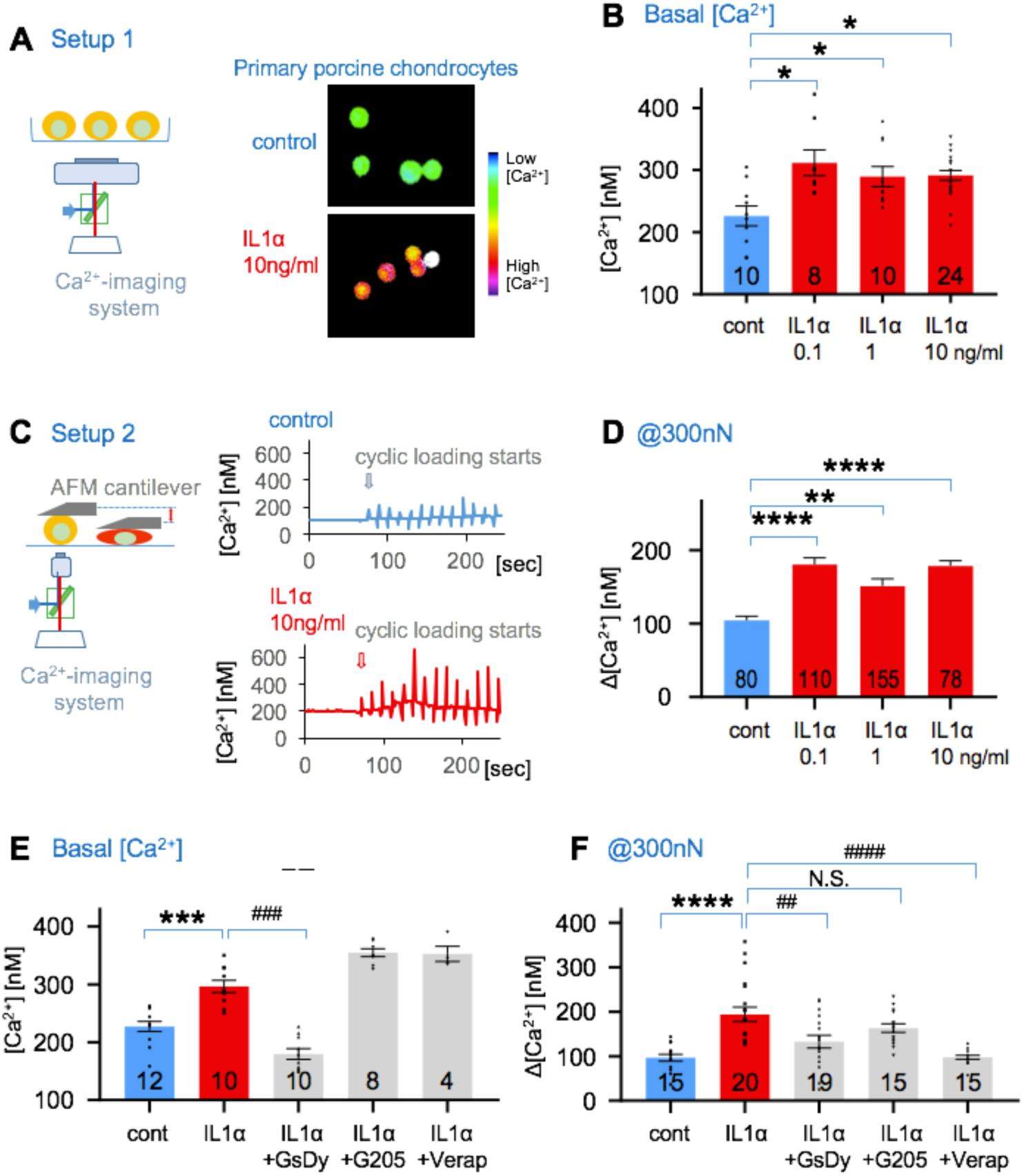
Pro-inflammatory IL-1α signaling alters Ca^2+^ dynamics in porcine articular chondrocytes. **(A)** (Left) schematic diagram of setup; (Right) representative ratiometric Ca^2+^ images of IL-1α-treated chondrocytes. **(B)** Resting cytosolic Ca^2+^ concentrations of primary articular chondrocytes are significantly increased, when exposing cells to IL-1α. Numbers indicate independent experiments, (bracketed numbers) indicate total numbers of cells measured. **(C)** (Left) schematic diagram of setup, AFM probe (flat, tip-less) compresses single cells cyclically every 10 sec; (Right) Ca^2+^ concentrations in response to cyclical compression. Loading starts at arrow-marked time-point. Representative Ca^2+^ transients are shown: (top) control, (bottom) IL-1α treated chondrocytes. **(D)** Mechanical compression-induced Ca^2+^ transients are significantly increased when exposing primary chondrocytes to IL-1α. Numbers indicate numbers of transients (increase in [Ca^2+^]i over resting [Ca^2+^]i) measured, number of cells stimulated were 16, 23, 31 and 16. **(E)** Resting [Ca^2+^]i of chondrocytes that were treated with IL-1α (1ng/ml), and inhibitors of Piezo (GsMTx4 2µM/dynasore 5µM), TRPV4 (GSK205 25µM) and voltage-gated Ca^2+^ channels (VGCC; verapamil 0.5µM). Resting [Ca^2+^]i that is significantly elevated by exposure to IL-1α is significantly reduced, below control values when inhibiting Piezo, but not decreased by inhibition of TRPV4 or VGCC. Numbers in bars indicate independent experiments, with numbers of cells examined 855, 457, 385, 387 and 153. **(F)** Experimental groups as in (E), but mechanical compression evoked Ca^2+^ dynamics were measured, as in (C-D). Note that significant IL-1α evoked Ca^2+^ increase is significantly attenuated when inhibiting Piezo, again no decrease when inhibiting TRPV4. Remarkably, there was a complete elimination of the IL-1α evoked Ca^2+^ increase when inhibiting VGCC, in striking contrast to effects of VGCC on resting [Ca^2+^]i (see E). Numbers indicate numbers of transients measured, as in (D), respective number of cells was 3, 4, 4, 3 and 3. Bars represent mean±S.E.M; for group comparison B, D, E-F: 1W-ANOVA, post-hoc Tukey test; * comparison IL-1α vs control, # comparison IL-1α plus treatments vs IL-1α. */# p < 0.05, **/## p<0.01, ***/### p<0.001, ****/#### p<0.0001 significantly different between groups.

We next examined the mechanism underlying this Ca^2+^-sensitization. Steady-state [Ca^2+^]_i_ was attenuated by GsMTx4 combined with dynasore, a combination previously shown to inhibit Piezo channels in articular chondrocytes (3, 4, 30). We observed no effect on [Ca^2+^]_i_ by inhibition of the chondrocyte mechanosensitive channel TRPV4 with selective inhibitor GSK205, or inhibition of voltage-gated Ca^2+^ channels (VGCC) with verapamil, also expressed in chondrocytes (Fig. 2E). Interestingly, the mechanical load-induced Ca^2+^-response was also dependent on Piezo, as shown by strong reduction of the increase in [Ca^2+^]_i_ by GsMTx4/dynasore (Fig. 2F). This finding indicated that increased expression of Piezo1 in IL-1α-mediated inflammation was responsible for baseline steady-state [Ca^2+^]_i_ increase as well as the mechanical load-evoked [Ca^2+^]_i_ increase. TRPV4 was not involved in either condition. We noted with great interest that VGCC were also key for the mechanical load-evoked [Ca^2+^]_i_ increase, in clear contrast to these channels’ lack of contribution to increased baseline steady-state [Ca^2+^]_i_. Therefore, the mechanical load-evoked [Ca^2+^]_i_ increase relies on both PIEZO1 and VGCC, potentially functioning as a Ca^2+^-amplification mechanism of directly-mechanically activated PIEZO1 channels, a finding consistent with our earlier characterization of PIEZO-mediated mechanotransduction in chondrocytes ^3^. This is not the case without mechanical load where increased baseline steady-state [Ca^2+^]_i_ relies solely on increased expression of PIEZO1 evoked by inflammatory IL-1α signaling, consistent with studies suggesting that low levels of PIEZO1 activation and inactivation are present at resting membrane tensions due in part to cytoskeletal forces (31-33). Taken together, increased Piezo1 expression evoked by IL-1α profoundly changes the basal function of articular chondrocytes by increasing the steady-state baseline [Ca^2+^]_i_ by >30%. Increased Piezo1 expression in turn makes these cells hypersensitive to cyclic mechanical loading with mildly injurious stimuli, resulting in additional Ca^2+^-influx. Additionally, VGCC function together with PIEZO1 only under direct deformational mechanical loading in the form of a mechanical stress amplification mechanisms, not at resting state.

### Piezo1 gain-of-function rarefies F-actin, makes chondrocytes vulnerable to mechanical injury

We next hypothesized that inflammatory signaling in articular chondrocytes might exert an effect on the cytoskeleton, the subcellular substrate of physical stiffness (modulus) of the cell, thus protecting or exposing chondrocytes to increased strain resulting from the thousands of cyclic mechanical loads each day that chondrocytes in weight-bearing joints undergo. We used phalloidin staining of porcine chondrocytes to visualize filamentous actin (F-actin), because force transmission through F-actin is an essential mechanotransduction process in chondrocytes (Fig. 3A) (34, 35). We found that F-actin was significantly reduced in IL-1α-exposed chondrocytes (Fig. 3B). If Piezo signaling was inhibited with GsMTx4/dynasore, F-actin was restored. This result indicates that Ca^2+^-influx through upregulated PIEZO1 leads to readily apparent changes of the chondrocytic cytoskeleton, namely a rarefication of F-actin. We did not find similar regulation of β-tubulin (Fig. 3C), nor did we see β-actin mRNA or total chondrocytic β-actin protein down-regulated in response to inflammatory IL-1α signaling (Suppl. Fig. 4). Thus, the F-actin rarefication in response to increased expression of PIEZO1 occurs at the level of intracellular protein processing and aggregation, not gene expression of β-actin. With its sub-plasmalemmal localization, F-actin provides structure and mechanical stability to the spherical chondrocytes upon compression. We next wanted to verify this concept in human OA cartilage and found that F-actin was also significantly reduced in OA articular chondrocytes, with F-actin rarefication appearing very similar to that observed in IL-1α-treated porcine articular chondrocytes (Fig. 3D).

**Fig. 3.**
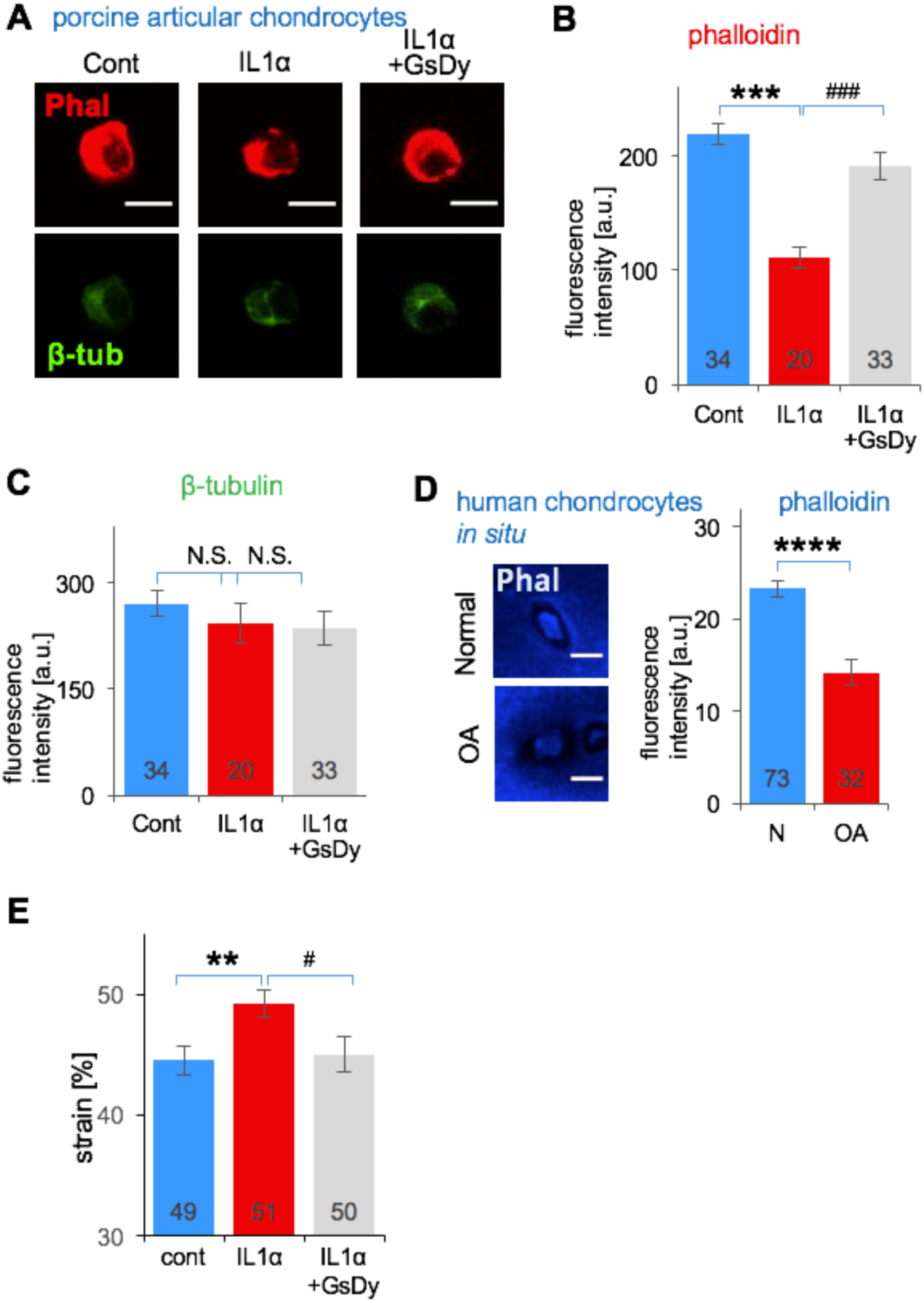
Rarefied F-actin cytoskeleton in IL-1α-exposed porcine articular chondrocytes and human OA cartilage. **(A)** Representative confocal fluorescent micrographs of porcine articular chondrocytes; cytoskeleton with F-actin (red) and β-tubulin (green) is visualized. **(B)** Quantitation of fluorescence intensity (arbitrary units (a.u.)) of F-actin (phalloidin labeling). IL-1α exposure significantly diminished F-actin, and Piezo-inhibition with GsMTx4/dynasore completely rescued the rarefication of the F-actin network. Numbers indicate number of cells analyzed. **(C)** Quantitation of fluorescence intensity of ß-tubulin labeling, note absence of significant changes. Numbers indicate number of cells analyzed. **(D)** Representative confocal fluorescent micrographs of human cartilage, healthy (normal) vs osteoarthritic (OA); cytoskeleton with F-actin (blue) is visualized. F-actin was significantly decreased in OA. Numbers indicate number of cells analyzed. **(E)** Strain on control, IL-1α-treated and IL-1α/GsMTx4/dynasore-treated porcine chondrocytes. IL-1α-treated cells (1ng/mL) have significantly higher strain than vehicle-treated cells. Note complete restitution to lower strain when inhibiting Piezo with GsMTx4/dynasore in IL-1α-treated chondrocytes. Numbers indicate number of cells analyzed, from 3 independent isolations of primary chondrocytes for each group. Bars represent mean±S.E.M; for group comparison B-C, E: 1W-ANOVA, post-hoc Tukey test; * comparison IL-1α vs control, # comparison IL-1α plus GsMtx4/Dyn vs IL-1α. */# p < 0.05, **/## p<0.01, ***/### p<0.001, significantly different between groups. Group comparison D: t-test; * p<0.05, **** p<0.0001, significantly different between groups.

We therefore investigated the influence of IL-1α on the mechanical properties and force-deformation behavior of porcine chondrocytes. Exposure to IL-1α led to significantly increased cellular deformation in response to the same magnitude of mechanical loading, a result of a decreased cellular modulus (Fig. 3E). This phenomenon translates to increased cellular mechanical strain, and potentially microtrauma upon compression. When inhibiting Piezo signaling with GsMTx4/dynasore, the cellular modulus and force-deformation properties reverted to control levels absent IL-1α. This finding, together with the PIEZO1-regulated F-actin polymerization, indicates that excess Ca^2+^ entering the chondrocyte via over-expressed PIEZO1, upon mechanical load, decreases cellular mechanical stiffness of chondrocytes due to rarefied F-actin in the cortical region of the cell. Decreased cellular rigidity will facilitate cellular mechanical microtrauma and thus increase likelihood of healthy chondrocytes transitioning into a degenerated OA phenotype. Importantly, both F-actin rarefication and decreased cellular rigidity in response to inflammatory IL-1α signaling were completely reversible when blocking Piezo signaling with GsMTx4/dynasore.

### IL-1α signaling in chondrocytes causing Piezo1 gain-of-function: from membrane to nucleus

Based on these findings, we next investigated signal transduction from membrane-bound IL-1 receptor (IL1R), via intracellular cytoplasmic signaling to nuclear transcriptional mechanisms that regulate the *PIEZO1* gene in articular chondrocytes in response to IL-1α. Since IL-1α will engage the IL-1 receptor type 1 (IL-1RI), known to be expressed by chondrocytes (10, 15), we assessed cellular signal transduction that upregulates expression of *PIEZO1* by using a candidate approach. We tested MAP-kinases, known to be down-stream of IL-1RI and downstream kinase, MKK3/6, namely MEK-ERK, JUN, and p38, also AKT-PI3K. Of these, only chemical inhibitors of p38-MAP kinase attenuated *PIEZO1* mRNA expression in porcine articular chondrocytes when exposed to IL-1α (Fig. 4A). Inhibiting the other kinases had no effect. In support, the phosphorylated isoform of p38 was significantly increased in response to IL-1α stimulation whereas its total protein abundance was unchanged (Fig. 4B). Phosphorylated p38 is known to traffic to the nucleus to regulate gene expression (36, 37). To identify transcription factors (TFs) that can upregulate *PIEZO1* in articular chondrocytes, we conducted a delimited TF screening experiment. We found 19 TFs to be upregulated in porcine articular chondrocytes in response to IL-1α, out of a total of 96 assessed TFs (Fig. 4C). We next analyzed *PIEZO1* regulatory DNA sequences from 2000 base-pairs upstream of the transcriptional start site to 500 bp down-stream of the transcriptional start site (TSS) in human, cow and chimpanzee. Using the MULAN DNA sequence analysis program, we generated predicted binding sites present in all three species tested (Suppl. Fig. 5). We further interrogated ATF2/CREBP1 and HNF4 because they identified from the 96 TF screen and were ≥4× up-regulated, they were present in all species’ proximal *PIEZO1* promoter in MULAN, and they can be reliably inhibited with well-established selective small molecule inhibitors (Fig. 4C, Suppl. Fig. 5). We selected HIF, NFAT and NF-κB as controls, expected not to regulate *PIEZO1* in response to IL-1α because they were not picked up by MULAN or the 96 TF screen. Of these TFs, in porcine articular chondrocytes, inhibition of ATF2/CREBP1 and HNF4 significantly and dose-dependently attenuated the increase of *PIEZO1* expression in response to IL-1α (Fig 4D). As expected, inhibition of HIF, NFAT and NFκB had no effect on *PIEZO1* expression. Of the TFs that we identified to enhance expression of *PIEZO1*, ATF2 and CREBP1 are family members known to bind to a consensus DNA binding site and to form protein-protein complexes that also include p38 (38, 39). HNF4 has also been demonstrated to be part of such multi-protein complexes by binding to p38 and to CREB (40, 41). Based on the MULAN bioinformatics analysis (Fig. 4E, Suppl. Fig. 5), we predicted direct binding of CREBP/ATF2 and HNF4 to the proximal *PIEZO1* promoter to regulate the *PIEZO1* gene, rather than indirect regulation through other transcriptional enhancer or de-repression mechanisms. We examined CREBP1-binding to the *PIEZO1* promoter by ChIP-qPCR, using DNA extracted from porcine articular chondrocytes after subjecting these cells to immuno-precipitation for CREBP1 (Fig. 4F). We amplified sequences from the predicted ATF2/CREBP1-binding site in the vicinity of the *PIEZO1* TSS. When immuno-precipitating with anti-CREBP1, we detected a robust >5x increase of *PIEZO1* promoter amplified DNA than with control antibody, indicating direct binding of CREBP1 to the probed site. Given the known interaction between CREBP1, ATF2 and HNF4 we regard it as highly likely that ATF2 and HNF4 also participate in a multi-protein enhancer complex that directly binds to the proximal *PIEZO1* promoter and upregulates *PIEZO1* gene expression.

**Fig. 4.**
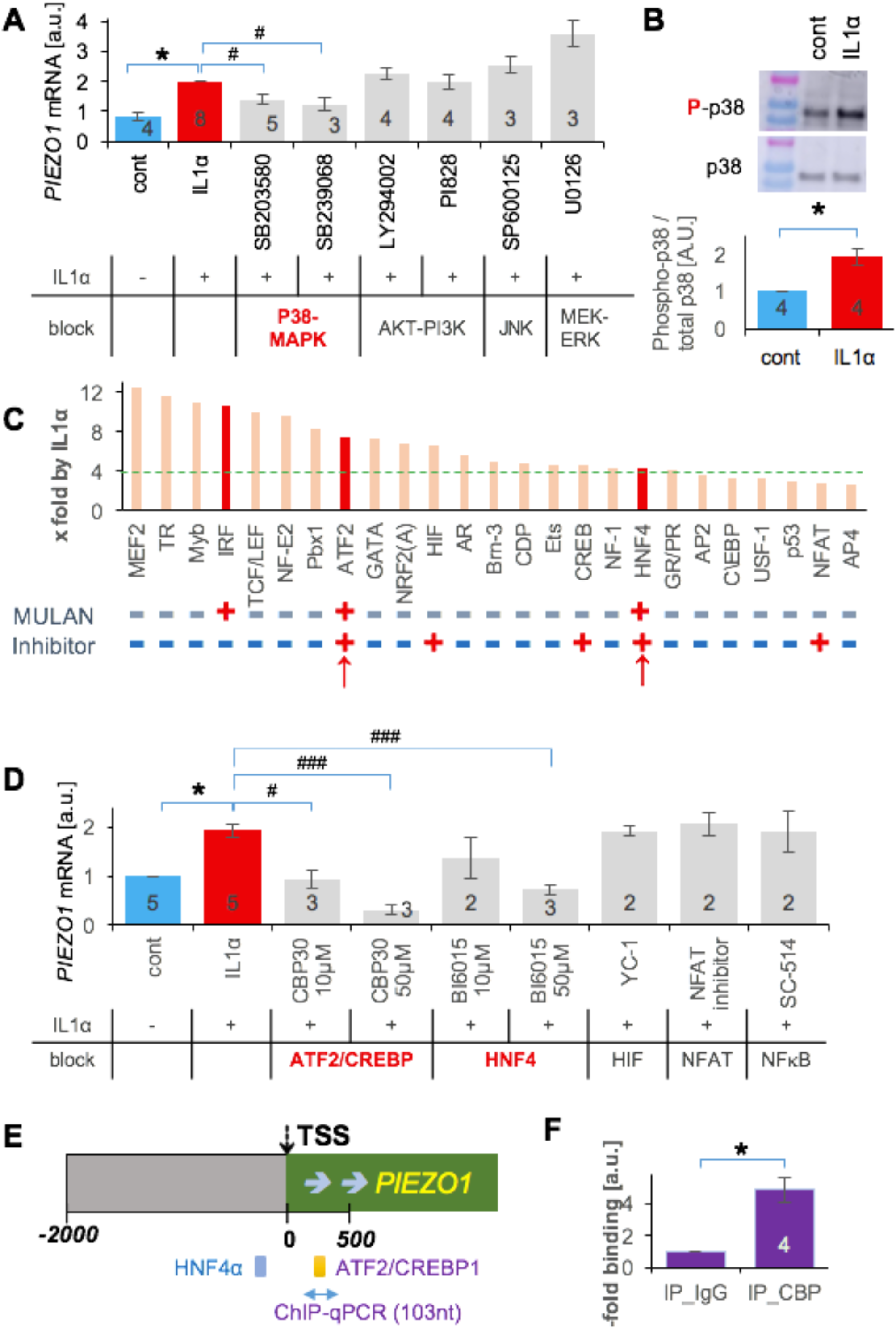
Identification of IL-1α-induced signal transduction that results in increased *PIEZO1* gene expression by delimited screening in porcine articular chondrocytes. **(A)** Inhibitors of p38 MAP-kinase significantly attenuated IL-1α-induced *PIEZO1*-mRNA increase (IL-1α 1ng/mL for 3 days, for all panels), but not inhibitors of MAP-kinases, JNK, MEK-ERK, and PI3K. **(B)** Western blot analysis of p38 phosphorylation indicates a significant increase of phospho-p38, total p38 does not change. **(C)** Starting with 96 transcription factors (TF), their activation was assessed in response to IL-1α treatment; increased x-fold over baseline, ranked, top-25 shown here. Note the green dotted line at 4x, with 19 of the top-25 TFs ≥4. Below the bar diagram, labeled “MULAN”, note binary score (+/-) whether the respective TF was found to have a predicted binding site in the proximal *PIEZO1* promoter (see E, Suppl. Fig. 5), using MULAN TF-binding prediction program. Indicated below, labeled “Inhibitor”, whether the respective TF can be inhibited with a well-established, selective small-molecule inhibitor compound. The two red arrows point toward the two TFs that fulfill all criteria, ATF2 and HNF4. **(D)** TFs that impact IL-1α-induced *PIEZO1*-mRNA increase were identified using selective compounds. HNF4 and ATF2/CREBP (the latter having the same DNA binding site as ATF2, known to form TF complexes) were confirmed as relevant, HIF, NFAT and NFkB were not found to be involved. Inhibitors of ATF2/CREBP1 and HNF4 significantly attenuated *PIEZO1*-mRNA increase. The dose-response relationship of ATF2/CREBP1 inhibitor, CBP30 showed a Pearson correlation coefficient = −0.85, p=0.0008, indicating a significant correlation. The respective metrics for HNF4 inhibitor BI6015 were Pearson correlation coefficient = −0.86, p=0.0014, also indicative of a signification correlation. **(E)** Schematic of predicted binding sites of HNF4, ATF2/CREBP1 TFs in the proximal *PIEZO1* promoter, as revealed by bio-informatics platform, MULAN. Please see also Suppl. Fig. 5. **(F)** Direct CREBP1 binding to the *PIEZO1* promoter was assessed by chromatin immunoprecipitation followed by qPCR. Note 5-fold increased abundance with IL-1α signaling vs control, indicative of direct CREBP1-binding to the *PIEZO1* proximal promoter site indicated in E (yellow bar, between 0 and +500 *PIEZO1* promoter). For A, B, D and F: numbers in bars indicate number of independent isolations of primary chondrocytes. Bars represent mean±S.E.M; for group comparison B, F: t-test, * p<0.05; for group comparisons A, D: 1W-ANOVA, post-hoc Tukey test; * comparison IL-1α vs control, # comparison IL-1α plus treatment vs IL-1α. */# p < 0.05

## Discussion

Taken together (Suppl. Fig. 6), we demonstrate a signaling mechanism from membrane to nucleus, from the IL-1α-IL-1RI complex, then via MKK3/6 to p38 MAP-kinase (as previously established), phospho-p38 to nuclear signaling of TF CREBP1 which binds *PIEZO1* regulatory DNA sequences together with TFs HNF4 and ATF2. These TFs then enhance *PIEZO1* gene expression. This signaling cascade leads to over-expression of Piezo1 in articular chondrocytes in response to inflammatory signaling. Over-expressed PIEZO1 channels in turn cause increased resting [Ca^2+^]_i_ and increased mechanical stress-evoked Ca^2+^ increase, indicative of mechanical hypersensitivity, “hyper-mechanotransduction” (42, 43). Increased resting [Ca^2+^]_i_, via PIEZO1, leads to rarefication of F-actin which decreases rigidity of the chondrocytes. This results in increased cellular deformation in response to mechanical loading. We recognize a detrimental feed-forward mechanism because increased deformation elevates the opening probability of mechanosensitive PIEZO channels, amplified by PIEZO1 being overexpressed. We speculate that the rarefied F-actin cytoskeleton enhances cytoplasmic availability of p38 for phosphorylation which happens after activation of IL-1RI, based on previous demonstration of the F-actin cytoskeleton influencing gene expression (44, 45).

In summary, we have identified a novel pathogenic feed-forward signaling and gene-regulatory mechanism in weight-bearing articular chondrocytes. This mechanism appears relevant for the pathogenesis of OA because it is a molecular mechanism linking inflammatory signaling with resulting hyper-mechanotransduction via upregulated *PIEZO1* expression and function. As one result of *PIEZO1* overexpression and resulting excess steady-state Ca^2+^, we found cytoskeletal changes of F-actin that facilitate cellular mechanotrauma, predisposing and contributing to chondrocyte dedifferentiation and degeneration as the foundation of the irreversible cartilage damage in “degenerative” OA. Breaking through this feed-forward mechanism therapeutically provides a rational target in the quest for OA disease-modifying therapies.

## Materials and Methods

### Porcine chondrocyte isolation and IL-1α treatment

Primary porcine articular chondrocytes were cultured as described in (3). These cells were exposed to the pro-inflammatory cytokine, IL-1α (at 0, 0.1, 1, 10 ng/ml) for 3 days) (28, 29).

### Measurement of steady-state intracellular Ca^2+^ concentration

To measure cytosolic [Ca^2+^] level, chondrocytes were loaded with Ca^2+^ sensitive dye, Fura-2-AM as described in (3).

### Measurement of compression-induced Ca^2+^ transients

Mechanically-induced Ca^2+^-influx of individual chondrocytes was measured using a custom-built AFM/Ca^2+^ setup consisting of an atomic force microscope (Bioscope; Veeco) and a ratiometric Ca^2+^ imaging microscope (Intracellular Imaging; Fig. 2C) as described in (3).

### mRNA expression in isolated chondrocytes by RT-qPCR

After treatments, chondrocytes’ total RNA was extracted using Trizol for RT-qPCR. PCR specificity was confirmed by gel electrophoresis and dissociation curve analysis. GAPDH or 18S were used as a housekeeping gene for normalization, and the gene-under-study mRNA level was quantified using the 2^-ΔΔCT^ method, as in (46).

### Immunocytochemistry

Porcine articular chondrocytes were cultured on glass coverslips, fixed with 4% paraformaldehyde at 4°C for 15 min, permeabilized with 2% Triton X-100, blocked with 5% donkey or goat serum for 30 min. Then cells were exposed to primary antibodies at 4dC overnight, then to fluorescent secondary antibody. Immunolabeling was visualized using a Zeiss 780 confocal microscope. Anti-Piezo1 (NBP1-78537, Novus), anti-p38 (#8690, Cell Signaling) and anti-phospho-p38 (#4511, Cell Signaling) were used. Chondrocytes were also labeled with fluorescent Phalloidin (phalloidin-CF568, Biotium) to visualize F-actin, or with BT7R-dylight488 (Thermofisher) to visualize ß-tubulin, then imaged.

### Immunohistochemistry

Paraffin-embedded tissue sections (15 µm) of human articular cartilage from healthy (n=5) and OA-positive (n=6) anonymized donors were used (Articular Engineering, IL). Sections were immunolabeled with Piezo1-antibody (NBP1-78537) followed by fluorescent secondary antibody (Alexa Fluor 594) and imaged. For F-actin staining, sections were labeled with fluorescent Phalloidin (phalloidin-CF350, Biotium) and imaged.

### Treatment with compounds and GsMTx4

Chondrocytes were treated with compounds for 1∼2hr before and during fura-2 loading: GsMTx4 peptide (2 μM, provided by Philip Gottlieb, SUNY Buffalo), dynasore (5 μM, Tocris), verapamil (0.5 μM, Sigma). To identify signal transduction mechanisms and transcription factors that regulate IL-1α-induced Piezo1 expression, inhibitors were co-incubated with IL-1α (1 ng/ml) for 3 days. Inhibitors used were SGC-CBP30 (10-50 μM, Cayman), BI6015 (10-50 μM, Cayman), YC-1 (10 μM, Cayman), NFAT inhibitor (10 μM, Cayman), SB203580 (10 μM, VWR), SB239063 (10 μM, Sigma), LY294002 (10 μM, VWR), PI828 (10 μM, Fisher), SP600125 (10 μM, VWR), U0126 (10 μM, VWR).

### Chondrocyte strain measurement

To measure the applied strain to the cells during loading with AFM, a costum-written MATLAB code was used. Briefly, both the contact point between the cell and the cantilever and the height of the cell were measured with code. The strain applied to the cells was then determined by the code by using these two measures.

### Discovery of transcription factors regulating the IL-1α-induced Piezo1 expression by Transcription Factor Array

Nuclear extracts of porcine chondrocytes were prepared using Nuclear Extraction Kit (SK-0001, Signosis), then next the Transcription factor (TF) Activation Profiling Plate Array II (FA-1002, Signosis, CA) was used to monitor 96 TFs simultaneously, according to the manufacturer’s protocol.

### Transcription factor-binding site analysis by bioinformatics

TF-binding sites within the *PIEZO1* promoter were analyzed and predicted using the comprehensive, web-based Evolutionary Conserved Region Browser (http://ecrbrowser.dcode.org). The promotor regions of *PIEZO1* [-2000nt to +500nt, transcription start site (TSS) defined as zero] in three different species were aligned, using high-quality genome sequences with a defined *PIEZO1* TSS: human (hg19 Chromosome 16: 88853372-88850872), cow (bostau6 Chromosome18: 14041407-14039769) and chimpanzee (pantro3 Chromosome16: 88511045-88508550). The MULAN TF-binding prediction algorhythm, contained within ecrbrowser, was run on these three species to identify candidate TF-binding sites.

### Chromatin immunoprecipitation (ChIP) assay

ChIP assays were performed using primary articular porcine chondrocyte nuclear extract, cell isolation from 4 individual pigs was conducted. The Magnify ChIP kit (ThermoFisher) was used according to manufacturer’s instructions and as described previously (46, 47).

Anti-CRBP1-immunoprecipitated DNA was purified to perform qPCR to determine the relative abundance of *PIEZO1* promoter DNA fragments. Conserved sequences of cow (bostau6 Chromosome18: 14041407-14039769, AC_000175.1) and pig (NC_010448.3) were aligned to generate primers for ChIP-qPCR.

### Statistical Analysis

Data are presented as mean±SEM. Student t-test or one-way ANOVA, post-hoc Tukey test for group-to-group comparisons, were used to determine the statistical significance. For correlation, Pearson’s correlation coefficient and its p-value were calculated. All statistical analysis was conducted using GraphPad Prism 8.2.

## Note

The authors declare not to have a conflict of interest.

## Acknowledgments

**General**: We thank Dr. Michelle Yeo (Duke University) for assistance with TF array tests, Dr. Jorg Grandl (Duke University) for donating the *PIEZO*1^-/-^ HEK293t cell line, for discussion of PIEZO channel activity and for careful read of the manuscript. Dr Andrey Bortsov (Duke University) provided advice with statistical testing.

**Funding:** This work was supported by NIH grants AR074240, AR073221, AG15768, AG46927, DE027454, P30 AR069655, P30 AR074992, and P30 AR073752, and the Arthritis Foundation.

## Author contributions

W. Lee carried out experiments and analyzed data. W. Lee, H. Leddy, A. McNulty isolated cells, R. Nims and A. Savadipour conducted strain measurements, W. Lee and F. Liu ran ChIP assays, W. Lee and Y. Chen carried out Western blots. W. Lee, F. Guilak and W. Liedtke directed the study and wrote the paper.

## Supplementary Material (Figures 1-6; Methods)

**Supplementary Fig. 1.**
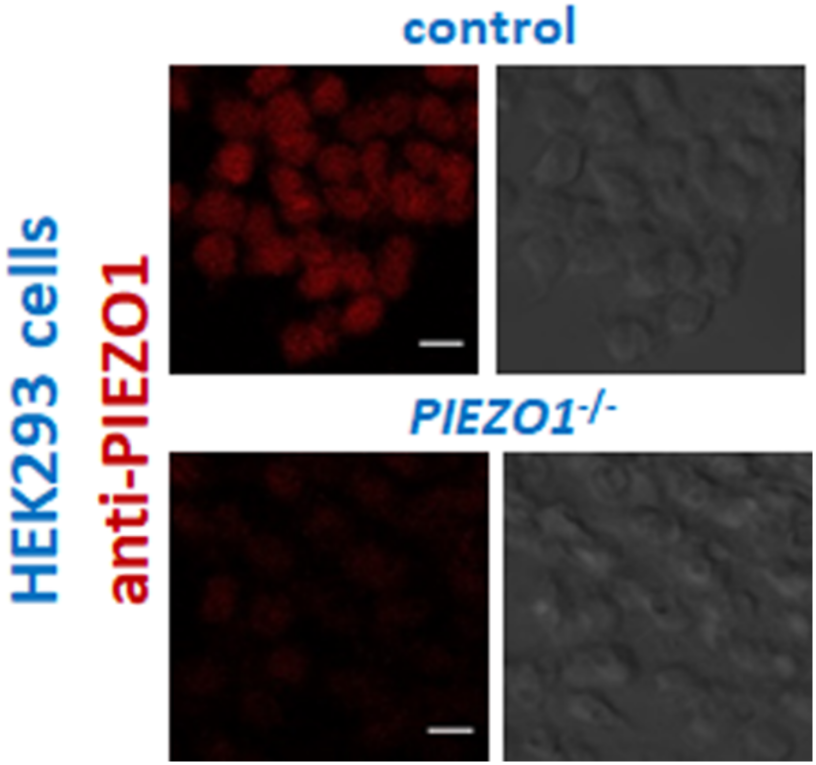
Antibody validation anti-PIEZO1 in HEK293 cells. Upper panels, wt HEK293t cells, showing a moderate level of PIEZO1 expression, left-hand side immunolabeling, right-hand side dark-field. Lower panel shows the respective findings in a genome-engineered HEK293t *PIEZO*^-/-^ cell line, indicating specificity of the used antibody in a human cell line.

**Supplementary Fig. 2.**
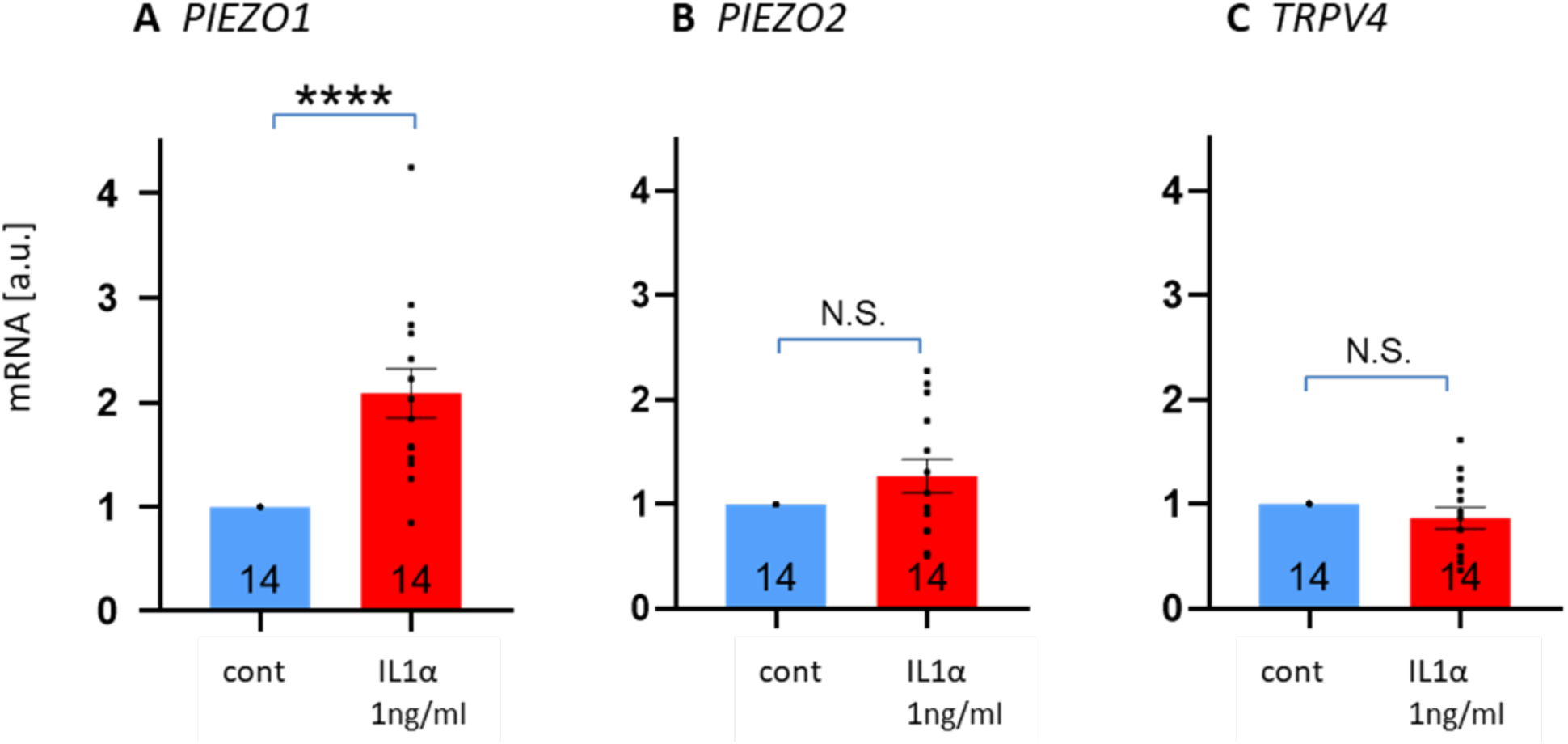
mRNA abundance for *PIEZO1, PIEZO2* and *TRPV4* in IL-1α-treated porcine articular chondrocytes. Numbers in bars indicate number of independent isolations of primary chondrocytes. Bars represent mean±S.E.M; for group comparison: t-test, p value indicated.

**Supplementary Fig. 3.**
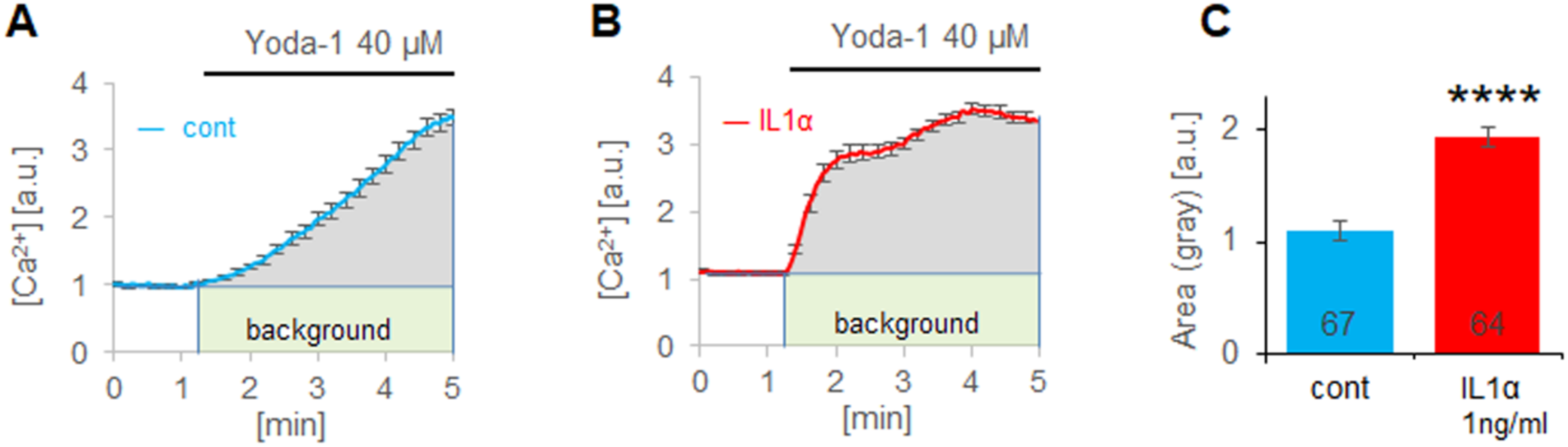
Yoda1-induced cytosolic [Ca^2+^] in chondrocytes. **(A-B)** Ca^2+^-transient area calculation as area-under-the-curve above ordinate=1 (=background). **(C)** Area-under-the-curve is significantly larger for IL-1α–treated chondrocytes as compared to control cells indicative of robustly increased Ca^2+^-influx via PIEZO1. Numbers in bars indicate number of individual cells imaged. Bars represent mean±S.E.M; for group comparison: t-test, ****p<0.0001.

**Supplementary Fig. 4.**
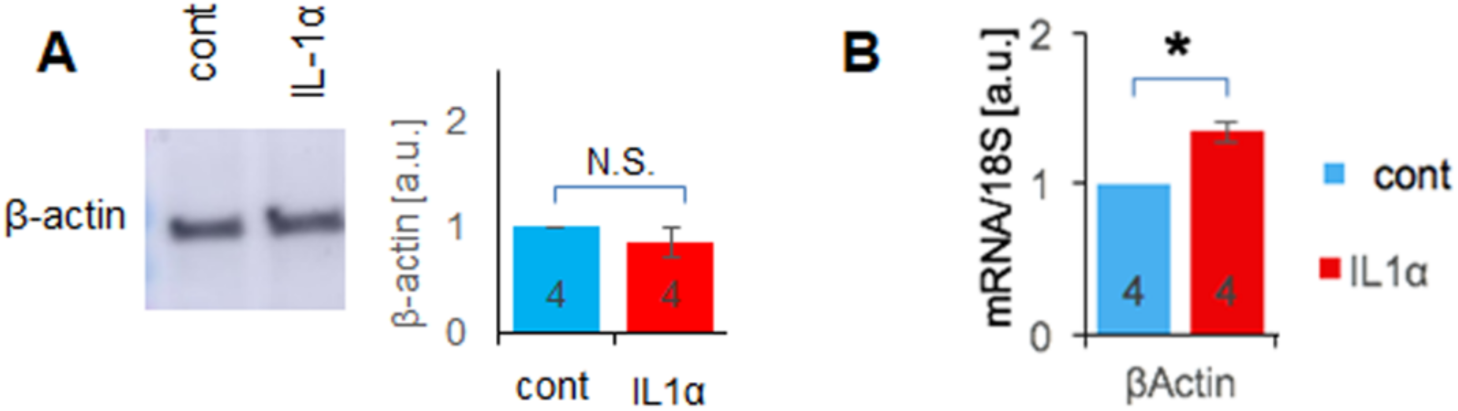
ß-actin expression in primary articular porcine chondrocytes. **(A)** Equal level of protein expression of ß-actin in porcine articular chondrocytes exposed to IL-1α or control. Quantitation of 4 independent experiments, densitometry of Western blotting bands, equal amount total protein loaded from primary chondrocyte cultures. **(B)** mRNA for ß-actin is not decreased in porcine articular chondrocytes exposed to IL-1α, it is rather elevated. Quantitation of 4 independent experiments, RT-qPCR, normalized for 18S RNA. * p<0.05, t-test. Numbers in bars indicate independent chondrocyte isolations.

**Supplementary Fig. 5.**
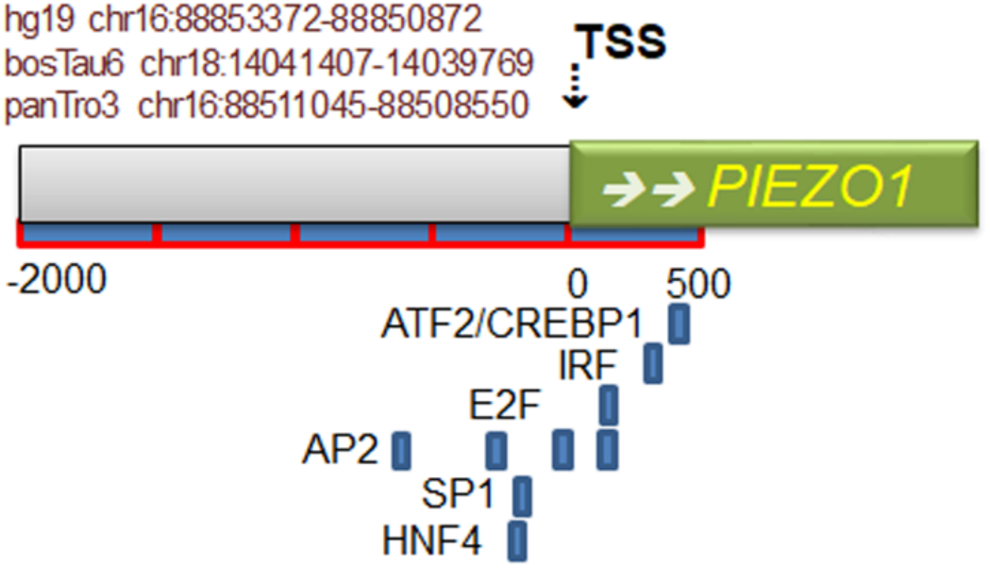
MULAN predicted TF-binding sites in the proximal *PIEZO1* promoter. The analysis is based on the genomic sequences of human, cow and chimpanzee. Only binding sites present in all three species were recorded. TSS – transcriptional start site.

**Supplementary Fig. 6.**
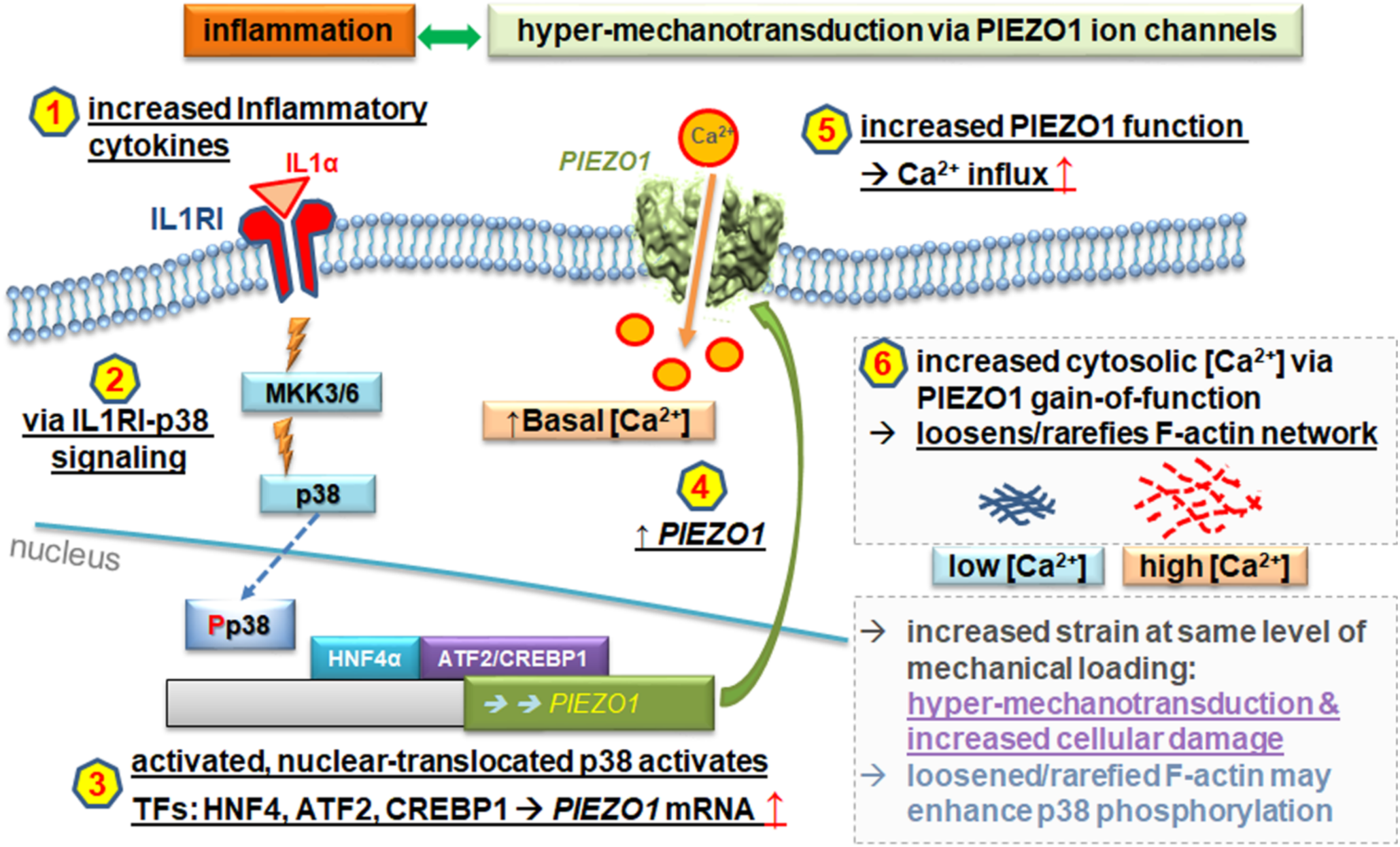
Feed-forward pathogenesis of OA relying on chondrocytic inflammatory signaling which results in Piezo1-gain-of-function. Schematic of our findings and proposed new OA pathogenetic mechanism. Signaling hubs and their consequences, upon activation as we demonstrate, are shown as a sequence 1-6. 1-6 in black letters is supported by our findings and known background. We interpret the result as “hyper-mechanotransduction & increased cellular damage”, in purple, and speculate, in blue, that a loosened/rarefied F-actin cytoskeleton may enhance p38 phosphorylation by making p38 more available for kinases.

## Supplementary Methods

### Porcine chondrocyte isolation and IL-1α treatment

Primary articular chondrocytes were harvested from articular cartilage of the femoral condyles of skeletally mature (2-3 year old) female pigs as described in (3). The isolated chondrocytes were seeded on 12-mm coverslips and tissue culture plates. Next day, chondrocytes were exposed to the pro-inflammatory cytokine, IL-1α, at 0, 0.1, 1, 10 ng/ml for 3 days (28, 29). Ca^2+^ imaging, mechanical stimulation experiments and RT-qPCR were performed 60∼72 hours after IL-1α treatment. Culture medium was changed every 2 days.

### Measurement of steady-state intracellular Ca^2+^ concentration

To measure cytosolic [Ca^2+^] level, chondrocytes on coverslips were loaded with Ca^2+^ sensitive Fura-2-AM dye (2 μM for 40 min; Invitrogen) and imaged using a ratiometric Ca^2+^ imaging platform (Intracellular Imaging, Setup 1 in Fig. 2A) as described in (3). Cytosolic [Ca^2+^] was determined using Incytim-2 software (Intracellular Imaging), based on 340/380 ratio and calibration of standard solutions, measuring [Ca^2+^]_i_ in individual chondrocytes.

### Measurement of compression-induced [Ca^2+^] transients

Mechanically-induced Ca^2+^-influx of individual chondrocytes was measured using a custom-built AFM/Ca^2+^ setup consisting of an atomic force microscope (Bioscope; Veeco) and a ratiometric Ca^2+^ imaging microscope (Intracellular Imaging; Fig. 2C) as described in (3). Briefly, chondrocytes on coverslips were loaded with Fura-2-AM and were compressed by a tip-less AFM probe with a force of 300 nN cyclically every 10 seconds for 2 min. Spring constants of the tip-less cantilever were 0.5–14 N/m (Novasan or Bruker Probes), the compression rate was 1– 2 μm/s, and the experiments were conducted at 37°C. The five maximum Ca^2+^-influx of each chondrocyte were determined ([Ca^2+^]_max_), and influx was determined as Δ[Ca^2+^]=[Ca^2+^]_max_ - [Ca^2+^]_i_ss_.

### mRNA expression in isolated chondrocytes by RT-qPCR

On the third day after cytokine or chemical treatments, chondrocytes were lysed and total RNA was extracted using Trizol, followed by RT-qPCR. PCR specificity was confirmed by gel electrophoresis and dissociation curve analysis. GAPDH or 18S were used as a housekeeping genes for normalization, and the gene-under-study mRNA levels were quantified using the 2^-ΔΔCT^ method, as in (46).

### Immunocytochemistry

Porcine articular chondrocytes and *PIEZO*1^-/-^ HEK293t cells were cultured on glass coverslips, fixed with 4% paraformaldehyde at 4°C for 15 min, permeabilized with 2% Triton X-100, blocked with 5% donkey or goat serum for 30 min. Then cells were exposed to primary antibodies at 4°C overnight, then to fluorescent secondary antibody. Immunolabeling was visualized using a Zeiss 780 confocal microscope. Anti-Piezo1 (NBP1-78537, Novus), anti-p38 (#8690, Cell Signaling) and anti-phospho-p38 (#4511, Cell Signaling) were used. Chondrocytes were also labeled with fluorescent Phalloidin (phalloidin-CF568, Biotium) to visualize F-actin, or with BT7R-dylight488 (Thermofisher) to visualize ß-tubulin, then imaged. A Zeiss780 confocal imaging system was used.

### Treatment with compounds and GsMTx4

Chondrocytes were treated with compounds for 1∼2hr during Fura-2A loading process: GsMTx4 peptide (2 μM, provided by Philip Gottlieb, SUNY Buffalo), dynasore (5 μM, Tocris), verapamil (0.5 μM, Sigma). To identify signal transduction mechanisms and transcription factors that regulate IL-1α-induced Piezo1 expression, inhibitors were co-incubated with IL-1α (1 ng/ml) for 3 days. Inhibitors used were SGC-CBP30 (10-50 μM, Cayman Chemical), BI6015 (10-50 μM, Cayman), YC-1 (10 μM, Cayman), NFAT inhibitor (10 μM, Cayman), SB203580 (10 μM, VWR), SB239063 (10 μM, Sigma Aldrich), LY294002 (10 μM, VWR), PI828 (10 μM, Fisher Scientific), SP600125 (10 μM, VWR), U0126 (10 μM, VWR).

### Chondrocyte strain measurement

To measure the applied strain to the cells during loading with AFM, a costume written MATLAB code was used. Briefly, both the contact point between the cell and the cantilever and the height of the cell were measured with code. At the end, using these two variables, the strain applied to the cells was determined by the code.

### Discovery of transcription factors regulating the IL-1α-induced Piezo1 expression by Transcription Factor Array

Nuclear extracts of porcine chondrocytes were prepared using Nuclear Extraction Kit (SK-0001, Signosis), then next the Transcription factor (TF) Activation Profiling Plate Array II (FA-1002, Signosis, CA) was used to monitor 96 TFs simultaneously, according to manufacturer protocol. Briefly, the nuclear extracts were incubated with biotin-labeled probes which were designed based on the consensus sequences of TF-binding sites. The TF-probe complexes were purified, then the bound probes separated from the complex. The detached probes were hybridized in 96 well plates where each well is specifically coated with complimentary sequences of the probes. The bound DNA probes are mixed with Streptavidin-HRP conjugates, and the luminescence was read in microplate luminometer. The luminescence of control and the IL-1α-treated chondrocytes were compared.

### Chromatin immunoprecipitation (ChIP) assay

Anti-CRBP1-immunoprecipitated DNA was purified to perform qPCR to determine the relative abundance of *PIEZO1* promoter DNA fragments. Conserved sequences of cow (bostau6 Chromosome18: 14041407-14039769, AC_000175.1) and pig (NC_010448.3) were aligned to generate primers for ChIP-qPCR. A 103 nucleotide amplicon was generated from the pig *PIEZO1* promoter: (CTCCGGATTAAACAGCTCCAGGCAGGAAGC-CCGCCTTCTCCCAGATTGGTCAGGAAGTCC<u>TGATGCAA</u>GTTTGCGCGTTTTCTTTCTCTC TCTCTCTCTCTTT; CREBP1-binding site is underlined). The sequences of the forward and backward primers are: For-CTCCGGATTAAACAGCTCCA and Rev-AAAGAGAGAGAGAGAGAGAAAGAAA.

